# Harmonizing Repeated Multi-Platform Optical and LiDAR Observations for Continental-Scale Terrestrial Carbon Cycle Data Assimilation

**DOI:** 10.64898/2026.10.30.763121

**Authors:** Dongchen Zhang, Qianyu Li, Shawn Serbin, Michael Dietze

## Abstract

Accurate monitoring of terrestrial carbon has attracted considerable attention due to the impacts of rapid climate change, disturbance, and land use on carbon storage and the growth of new global carbon markets and reporting requirements. While uncertainties in C estimates remain nontrivial, crucial opportunities exist to reduce uncertainty by rigorously harmonizing models and monitoring through data assimilation (DA). LiDAR has received considerable attention for its potential to inform carbon monitoring, and while there have been a number of papers on how to initialize models using LiDAR data, it has yet to be integrated into DA systems. In this study, we integrate GEDI LiDAR into an established DA system that already assimilates optical observations of aboveground biomass (AGB) and leaf area index (LAI), microwave soil moisture (SM), and ground-based soil organic carbon (SOC) to constrain C cycling processes across the entire North American continent. Notably, this is the first time both optical and LiDAR AGB products have been iteratively harmonized within a DA system. Results show that by assimilating both optical and LiDAR observations, we reduced the uncertainty in woody biomass by 63.7% relative to unconstrained model simulations and 18% relative to our previous DA system, and achieved an average RMSE reduction of 38.3 kg/m2 across five AGB products. Importantly, LiDAR reduces DA biases in high AGB regions where optical AGB estimates were exhibiting saturation. Analysis further identified the leading factors driving spatial error patterns within the DA system to be data quality, cloud coverage, and elevation. Overall, by bridging the spatial and temporal limitations of individual satellite products, this approach offers a practical, generalized workflow for high-resolution terrestrial carbon MRV in complex, dynamic ecosystems.

## 1. Introduction

Terrestrial ecosystems represent one of the most dynamic yet uncertain components of the global carbon cycle, serving as crucial buffers against rising atmospheric carbon dioxide while simultaneously undergoing rapid shifts driven by climate change, natural disturbances, and land-use change (Li et al., 2026, Arora & Boer, 2010; Harris et al., 2016). Accurately monitoring these terrestrial carbon stocks and fluxes has become paramount—not only to advance fundamental ecosystem science, but also to satisfy emerging reporting mandates and support transparency within expanding global carbon markets. North America (NA) exemplifies these challenges; despite being one of the most intensely monitored regions on Earth, shifting disturbance regimes and land management practices have rendered its status as a net carbon sink increasingly variable (Hayes et al., 2018). While substantial uncertainties persist in regional carbon estimates, recent advances in harmonizing process-based models with diverse observational streams through state data assimilation (DA) offer a pathway to reduce prediction uncertainties, close critical accounting gaps, and deliver robust carbon monitoring, reporting, and verification (MRV) systems.

In our previous work, we successfully integrated multiple remote sensing data sources with process-based models using DA (Zhang et al., 2026a-c). However, due to their physical limitations, each sensor reveals only partial information about an ecosystem’s structure, function, and dynamics, depending on its specific wavelength range, modality, or both, with associated pros and cons. For example, in our previous North American DA study (Zhang et al., 2026b), assimilating optical remote sensing observations could lead to systematic biases (underestimation) for high-AGB forests, where spectral reflectances are saturating and no longer sensitive to changes in AGB increment (Zhao et al., 2016; Sa et al., 2024). Similarly, places with high cloud coverage experience increased uncertainty in optical remote sensing estimates (Zhai et al., 2018). In general, while optical remote sensing has provided extensive measurements globally for decades, these observations are indirect measures of aboveground biomass and rely heavily on climate and ecosystem conditions (Ma et al., 2024).

Light Detection and Ranging (LiDAR), on the other hand, advances the estimation of vegetation biomass by overcoming the saturation problem and by providing a direct measure of 3D structure. Therefore, LiDAR has been widely used to estimate vegetation biomass, resulting in improved biomass estimate accuracy (Ni-Meister et al., 2022; Thapa et al., 2025). However, compared with optical remote sensing, LiDAR measurements (e.g., GEDI L4A footprints, Patterson et al., 2019; ICESat series, Duncanson et al, 2026) are generally sparser and have limited spatiotemporal coverage, making it challenging to generate gridded annual estimates of biomass, with most products providing either a single snapshot or time-integrated products where a single image averages over multiple years. Such products lack temporal continuity and ignore the dynamic recovery of specific ecosystems through time series (Vallet & Mouillo, 2026). By contrast, DA approaches excel at integrating sparse data, as they can rely on process model projections, other data constraints, and spatial covariances between measured and unmeasured locations to estimate C pools for the times and locations with missing data. However, while there is a well-developed literature on initializing models using LiDAR (Ma et al., 2023), to the best of our knowledge, LiDAR data have not yet been integrated into land DA systems.

Therefore, neither LiDAR nor optical remote sensing can perfectly represent both 3-D structure and optical characteristics with complete spatiotemporal coverage. Relying on a single platform brings a trade-off between coverage and accuracy, making it crucial to harmonize both optical and LiDAR observations within a unified DA framework. By leveraging the complementary strengths of both data products, such an approach iteratively constrains model states to deliver spatially continuous, structurally accurate C reanalyses. To date, many studies have combined multi-source remote sensing products for accurate biomass estimation (e.g., GEDI and Sentinel, Zurqani, 2025; Hernández-Martínez et al., 2025; airborne LiDAR and Hyperspectral, Ferreira et al., 2024). Yet most studies that attempt to harmonize optical and LiDAR data typically rely on static parameter calibration or single-time optimizations. However, these methods fail to deliver fully constrained, spatiotemporal state estimates with complete uncertainty quantification. Thus, establishing an iterative DA system using both optical and LiDAR platforms is essential to close the gap.

In this study, we extended our previous North American DA work (Zhang et al., 2026b) to develop an adaptive GEDI aggregation algorithm and updated our Tobit-Gamma Ensemble Filter to harmonize Landsat and GEDI data. To understand the patterns of improvements in AGB estimations relative to unconstrained (no DA) model simulations and to the results from our previous study (assimilating optical remote sensing only). This study is going to answer the following questions:

**Q1:** What are the spatiotemporal patterns for the residual and predictive errors relative to the baseline model and to the results of the previous North American DA work?

**Q2:** What are the driving factors that affect both the residual and predictive errors?

## 2. Methods

### 2.1. Study Area

This study examines the North American (NA) continent (-179° W to -20° W; 7° N to 85° N) at a 1-km spatial resolution with annual time steps from 2012 to 2024, covering more than 21 million km2 of vegetated land, including Central America, the Caribbean, Mexico, the USA mainland, Hawaii, Alaska, and Canada. Using the site-selection workflow described in Supplement 1, we selected 8,000 locations representing eight land-cover classes (Figure 1) based on the MODIS PFT land-cover classifications and 15 level-1 EPA ecoregions (Omernik, 1995). To select locations with the most GEDI coverage, we used weighted sampling based on the number of high-quality GEDI tiles. Throughout the GEDI period, 4,316 locations are covered by GEDI L4A footprints, with 3,752 locations having data over multiple time points.

**Figure 1.**
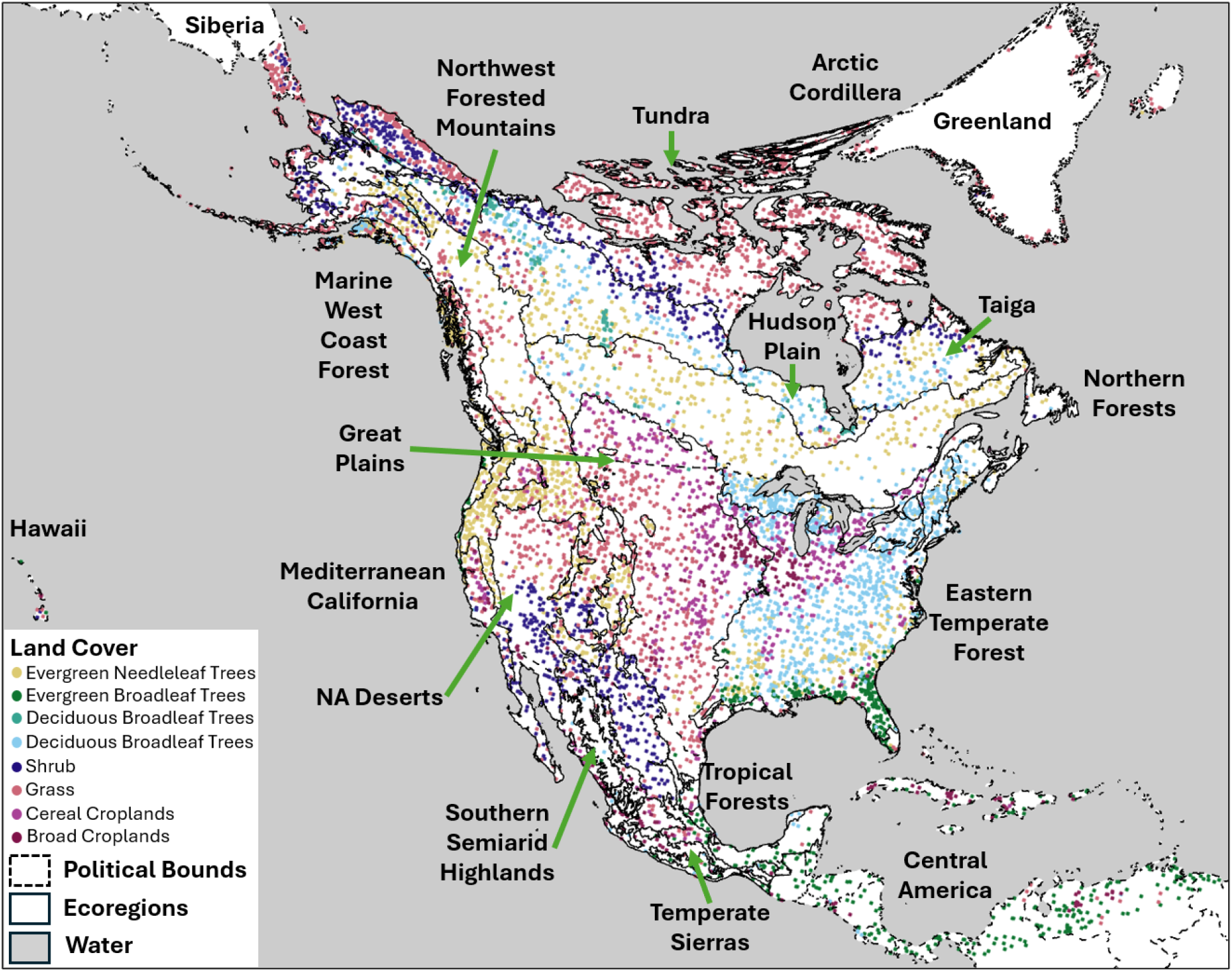
Pre-selected 8,000 locations based on our site-selection algorithm. The map is in an Albers projection to preserve the area across latitude.

### 2.2. GEDI Aggregation

After the site selection, we will need to prepare data inputs for each location. In addition to the observations used in the previous North American DA studies (LandTrendr AGB, MODIS LAI, SMAP SM, and SoilGrids SOC, Zhang et al., 2026a), this section introduces the algorithm for aggregating the new GEDI L4A AGB footprints. GEDI is a high-resolution (25m) spaceborne LiDAR instrument installed on the International Space Station that maps the 3-D structure of Earth’s surface and forests from 2019 to the present. In this study, we aggregated GEDI-L4A footprints using an annualized implementation of the GEDI Level 4B algorithm (Patterson et al., 2019) to produce gridded AGB estimates within a predefined spatiotemporal window. The GEDI-L4A AGB estimates have a 25 m resolution and were derived using linear regressions between the GEDI Level 2A relative height metrics and AGB field measurements across five PFTs and seven global regions (Dubayah et al., 2020). Next, we set up a series of one-year windows, each centered on July 15th, and a 1 km spatial window to match the MODIS grid used in the DA system. For example, in Figure 2 we can see five tracks representing standalone GEDI Level 4A footprints from descending and ascending orbits. Based on the GEDI-L4B algorithm, the aggregated AGB mean will be calculated based on the following equation:

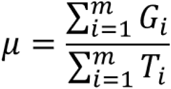

Where *m* stands for the total number of tracks within the boundary, *G*_*i*_ represents the total AGB for the *i*^*th*^ tracks, and *T*_*i*_ represents the total number of footprints for the *i*^*th*^ tracks. The aggregation of uncertainty includes two parts: the sample error and the parameter error, representing 1) sample sparsity (proportion of available GEDI tracks to the total possible number of non-overlapping tracks within a grid cell, which has 40 x 40 = 1600 25-m pixels in total) and 2) parameter error during the linear regressions, respectively. As shown in the following equation, the total uncertainty includes both the sample and parameter uncertainties:

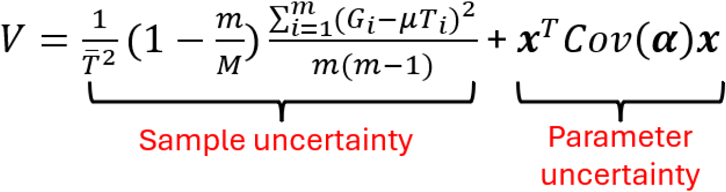

Where 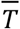 is the mean *T*_*i*_ across m GEDI tracks, *M* is the total number of possible tracks within the 1 Km cell, *x* is the vector of GEDI relative height metrics. Note that, in the equation, *x*_1_ equals 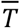 for mathematical convenience, where a Taylor expansion was used to approximate the value of the parameter uncertainty (Patterson et al., 2019). *Cov*(α) is the covariance matrix for the model parameters. In reality, because *m* << *M*, the above equation simplifies to:

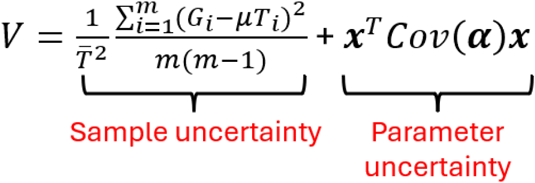

**Figure 2.**
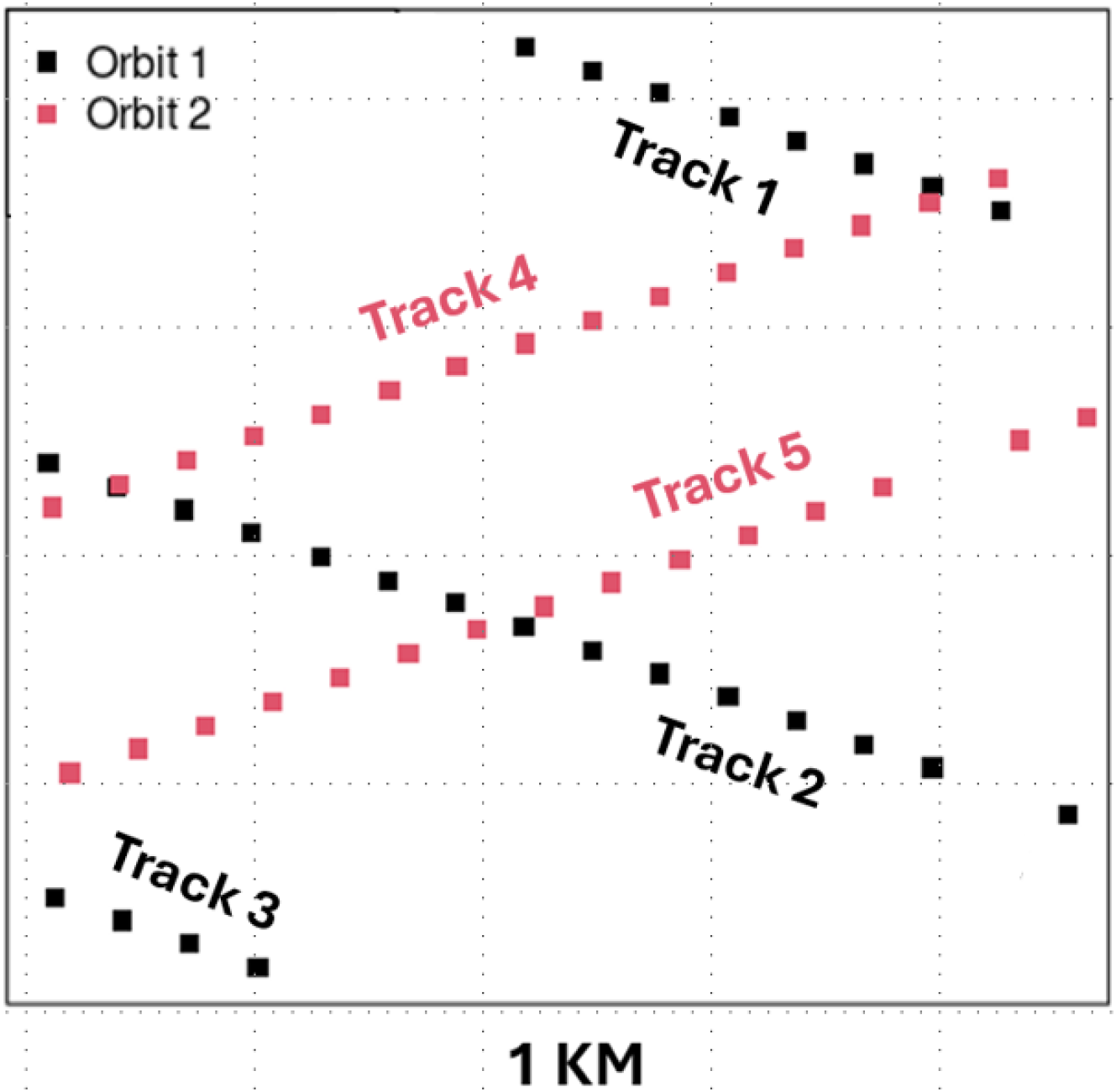
Diagram of GEDI footprints used to generate an area estimate of AGB within a 1km pixel. A GEDI track (m) represents a distinct set of GEDI footprints derived from a single orbit track crossing the 1 km×1 km target area during the designated annual observation period. The black and red tracks represent GEDI Level 4A footprints with descending and ascending orbits, respectively.

### 2.3. State Data Assimilation Workflow

In this study, we extended our previous Tobit Gamma Ensemble Filter (TGEnF; Zhang et al., 2026a) to account for multiple data streams from different platforms (e.g., optical and LiDAR remote sensing) constraining the same underlying state variable.

Our overall DA framework is shown in Figure 3. We selected the process-based model, SIPNET (https://github.com/PecanProject/sipnet, Braswell et al., 2005, Supplement 3), based on its low computational cost and its simplified process representations of coupled carbon and water cycling, which facilitate interpretability and data assimilation. For each of our 8000 sites, we ran 100 ensemble predictions of the SIPNET model, sampling across the uncertainty in the initial conditions (ICs, Supplement 4), calibrated parameters (Dokoohaki et al., 2022; Fer et al., 2018, 2021), and ERA5 meteorological drivers, each lasting 1 year from 2012 to 2024. Next, we assimilated the observations of major carbon and water pools (LandTrendr AGB, MODIS LAI, SMAP SM, and SoilGrids SOC, Table S3) into the ensemble forecasts using the TGEnF, harmonizing model simulations with multiple data streams and generating our reanalysis estimate of the C budget for that year. These outputs also serve as the ICs for the next round of one-year SIPNET forecasts. After the second year, we begin training a machine learning model to predict the previous forecast residuals (i.e., the forecast minus the observation) as a function of environmental covariates (Supplement 4), with the goals of (1) correcting model biases within the assimilation and (2) improving our understanding of the drivers of model bias. The bias correction is then applied iteratively until 2024, subtracting the predicted residuals from each modeled variable before feeding these predictions into the DA analysis step (Supplement 6).

**Figure 3.**
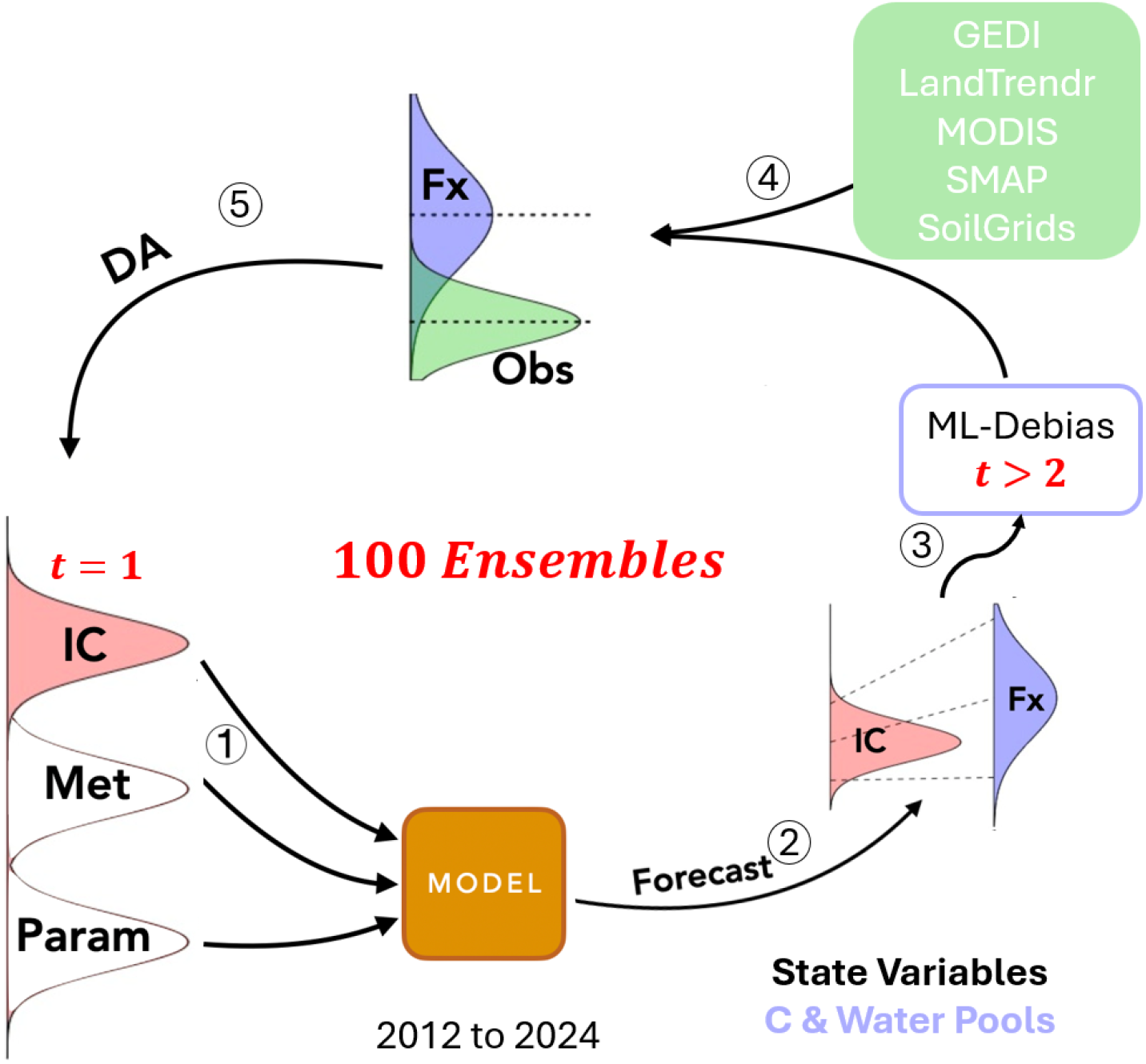
Workflow of the DA framework. (1) Sample ensemble members from the distribution of data inputs, including initial conditions, ERA5 meteorology drivers, and calibrated parameters. (2) Run an ensemble of the SIPNET model to produce forecasts (Fx) of key ecological states. (3) Apply the ML-Debias algorithm when t is greater than 2. (4) As new data become available for each timestep, (5) assimilate the four data streams (AGB, LAI, soil moisture, and soil carbon) using the MoTGEnF algorithm, producing harmonized estimates of the carbon and water pools that can be used as ICs for the next model forecast.

### 2.4. Experiments and Analysis

To assess the impact of assimilating GEDI AGB, we designed three experiments: the full-run assimilating GEDI and LandTrendr AGB, MODIS LAI, SMAP SM, and SoilGrids SOC; our previous North American DA experiments that constrained AGB with the LandTrendr product only (Zhang et al., 2026a); and a baseline run assimilating everything except AGB. This baseline experiment was chosen to exclude the beneficial impacts of indirect constraints (Zhang et al., 2026b), such that an unobserved but correlated variable or process (e.g., AGB in this study) will be inferred by other observations (MODIS LAI, SMAP SM, or SoilGrids SOC).

To address question 1 (Q1) about the following accuracy assessments focus on both residual and predictive errors of our AGB reanalysis (ensemble mean), we validated our AGB analysis against multiple AGB datasets (LandTrendr, Hooper & Kennedy, 2018; GEDI Level 4A and 4B, Dubayah et al., 2022; ICESat-2, Duncanson et al., 2023; FIA BIGMAP, Wilson et al., 2018; Table S2), while predictive errors are assessed using the ensemble standard deviation.

In comparing the full-run, previous DA run, and baseline-run (Q1), we assessed error patterns across both time and space. For the temporal patterns, we calculated the RMSE, R2, and relative changes between the full-run and baseline-run on an annual basis, averaging over the spatial domain of the corresponding observations. To assess spatial patterns, we calculated pixel-wise accuracy against the GEDI Level-4B dataset and uncertainty based on averaged AGB estimates from 2019 to 2024 to match the GEDI L4B period, as well as relative changes between the full-run and baseline-run. Note that, because GEDI L4B, FIA BIGMAP, and ICESat-2 are snapshots, we calculated the predictive skills of AGB within the temporal window for each product specifically (2019-2024 for GEDI, 2014-2018 for FIA BIGMAP, and 2019-2021 for ICESat-2).

To illustrate the improvements between the full run and the previous DA run (Q1), we visualized a series of scatter plots between our AGB reanalysis and the GEDI L4B observations, with R2 values across deciduous and evergreen forest types and latitude domains split by 30 and 50 latitudes, respectively. To assess the uncertainty reductions (Q2) from the full run relative to the baseline model, we calculated the averaged uncertainties across locations that were selected based on the site selection algorithm within the GEDI boundary (<52 latitude) for both experiments.

To understand the leading factors that affect both the residual and predictive errors (Q3), we hypothesized that the predictive skills will be strongly affected by 1) the number of GEDI measurements, as it directly controls the mean and uncertainty of aggregated GEDI AGB values; 2) elevation, as the topographical conditions are often challenging for accurate remote sensing estimates; 3) the number of annual cloudy days, as it can directly reduce the data quality of remote sensing observations because remote sensing products, especially the optical and LiDAR platforms, are highly vulnerable to dense clouds. Next, we visualized correlations between the number of unique GEDI tracks (10.3334/ORNLDAAC/2299) and residual and predictive errors in 2024 across the pre-selected locations below 52 latitude. We then used a Bivariate Local Moran’s I Analysis (Anselin et al., 2002) to assess the spatial correlation between error and each of 3 explanatory variables: number of GEDI tracks, elevation (U.S. Geological Survey, 2017), and annual cloudy days (Hall & Riggs, 2021). The Moran’s I analysis first applies a 12×12 moving window (144 pixels) and calculates the statistical correlations (p-value < 0.01) between the AGB predictive skills and the driving factors. There will be five clusters in the spatial correlation plots, including: 1) not significant (p-value > 0.01, grey); high accuracy/precision Vs. high driver values (the number of GEDI measurements, annual cloudy days, or elevation) (dark red); high accuracy/precision Vs. low driver values (light red), low accuracy/precision Vs. low driver values (dark blue), and low accuracy/precision Vs. high driver values (light blue) correlations.

## 3. Results

### 3.1. Predictive Skills and Uncertainty Assessments and Comparisons

As we can see from Figures 4a and 4b, assimilating LandTrendr AGB leads to increased predictive skills ( *R*^2^ from 0.91 to 0.99; RMSE from 1.39 to 0.48 *kg*/*m*^2^) between 2012 and 2017 compared to the LandTrendr observation, and a slight reduction in predictive skills ( *R*^2^ from 0.99 to 0.98; RMSE from 0.48 to 1.04 *kg*/*m*^2^) in 2018 due to data discontinuity when the LandTrendr AGB was reproduced using new Landsat time series, where LandTrendr shows a substantially significant reduction in AGB in 2018 compared to that of 2017 (Supplement 7). FIA validations show consistent accuracies from 2014 to 2018 (*R*^2^ of ∼0.73 and RMSE of ∼3.6 *kg*/*m*^2^). Similarly, the ICESat-2 validation shows a slight decrease in R2 (0.61 to 0.60) and consistent RMSE (1.78 to 1.79 *kg*/*m*^2^) from 2019 to 2021. Interestingly, assimilating both LandTrendr and GEDI observations leads to a continuous reduction in predictive skills (*R*^2^ from 0.98 to 0.85; RMSE from 1.04 to 2.28 *kg*/*m*^2^) for LandTrendr validation from 2018 to 2022. The GEDI-L4A validation shows a continuous increase in predictive skills for the full assimilation (*R*^2^ from 0.64 to 0.86; RMSE from 5.61 to 2.69 *kg*/*m*^2^) from 2019 to 2024. However, because GEDI was turned off from March 2023 to April 2024, the number of available GEDI tiles fell notably that year, reducing assimilated GEDI observations from 2939 to 194. This resulted in a slight increase in LandTrendr predictive skills (*R*^2^ from 0.85 to 0.92; RMSE from 2.28 to 1.63 *kg*/*m*^2^) and a slight decrease in *R*^2^ for GEDI-L4A validation (from 0.77 to 0.74), but a continuous decrease in the corresponding RMSE (from 3.59 to 2.92 *kg*/*m*^2^). The trend in GEDI-L4B *R*^2^ shows a slower increase (from 0.75 to 0.80) than the GEDI-L4A predictive skills trend, while the RMSEs remain nearly constant from 2019 to 2024.

**Figure 4.**
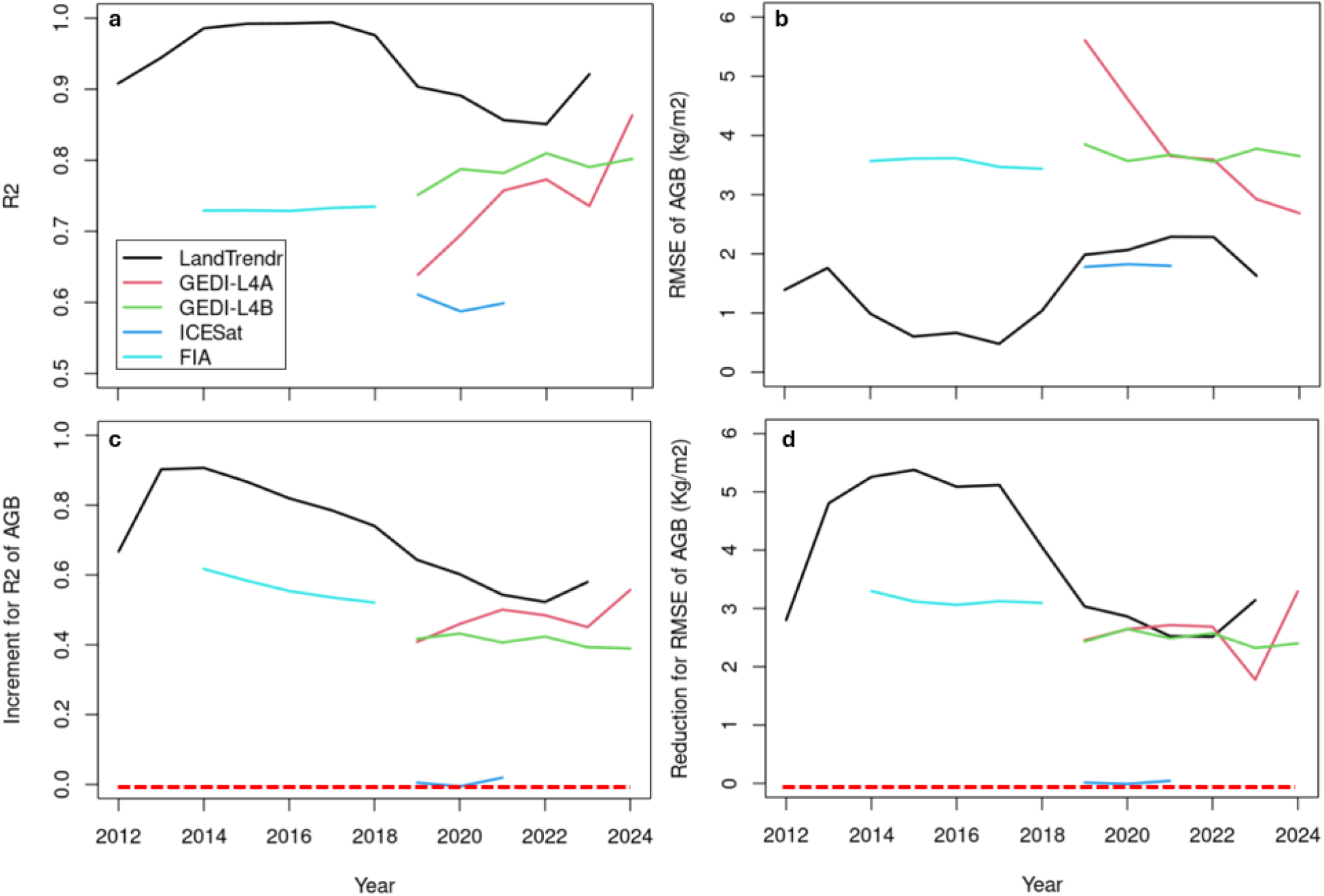
The temporal trends of the full run R^2^ and RMSE (a and b) and the relative changes in each between the full-run and baseline-run (c and d). Each line represents the R2 and RMSE of the full-run validations against different observations for the time period and spatial domain for which data were available. LandTrendr and GEDI-L4A were assimilated, so they are in the sample; GEDI-L4B is a time-averaged product and is not independent of L4A; ICESat and FIA are independent validations. (Note that the y=0 red dashed line represents the baseline model predictive skills).

Comparing the predictive skills of the full-run and baseline-run, Figure 4c shows that assimilating LandTrendr + GEDI AGB leads to *R*^2^ improvements, relative to the baseline model results, from 0.67 to 0.58 between 2012 and 2024. The improvement skill against the FIA BIGMAP showed a similar temporal trend, with an initial *R*^2^ improvement of 0.62 in 2014 and a slight decrease in *R*^2^ improvement over time, ending at 0.52 in 2018. The trend in *R*^2^ improvement against GEDI-L4A is similar to that of LandTrendr, while the reduction in GEDI-L4B *R*^2^ is slightly lower. Because of the GEDI pause in 2023, the improvements in both predictive skills drop relative to those in 2022. The ICESat predictive skills are similar between the full-run and baseline-run because the ICESat data product spans a latitudinal range (43.7°N to 78.5N) that is largely above the GEDI boundary (51.6°N latitude), resulting in a baseline-to-baseline comparison over much of the region. Figure 4d assesses improvements in RMSE and shows very similar patterns to Figure 4c.

Comparing the full-run DA results with the LandTrendr DA results, Figure 5 shows that assimilating GEDI consistently improves the predictive skill of AGB. Specifically, there are noteworthy improvements in AGB *R*^2^ values between 30 and 50 latitude (increases of 0.12 and 0.19 for deciduous and evergreen forests, respectively), where LandTrendr was widely available, as well as below 30 latitude (increases of 0.32 and 0.10 for deciduous and evergreen forests, respectively), where previously AGB was largely unconstrained.

**Figure 5.**
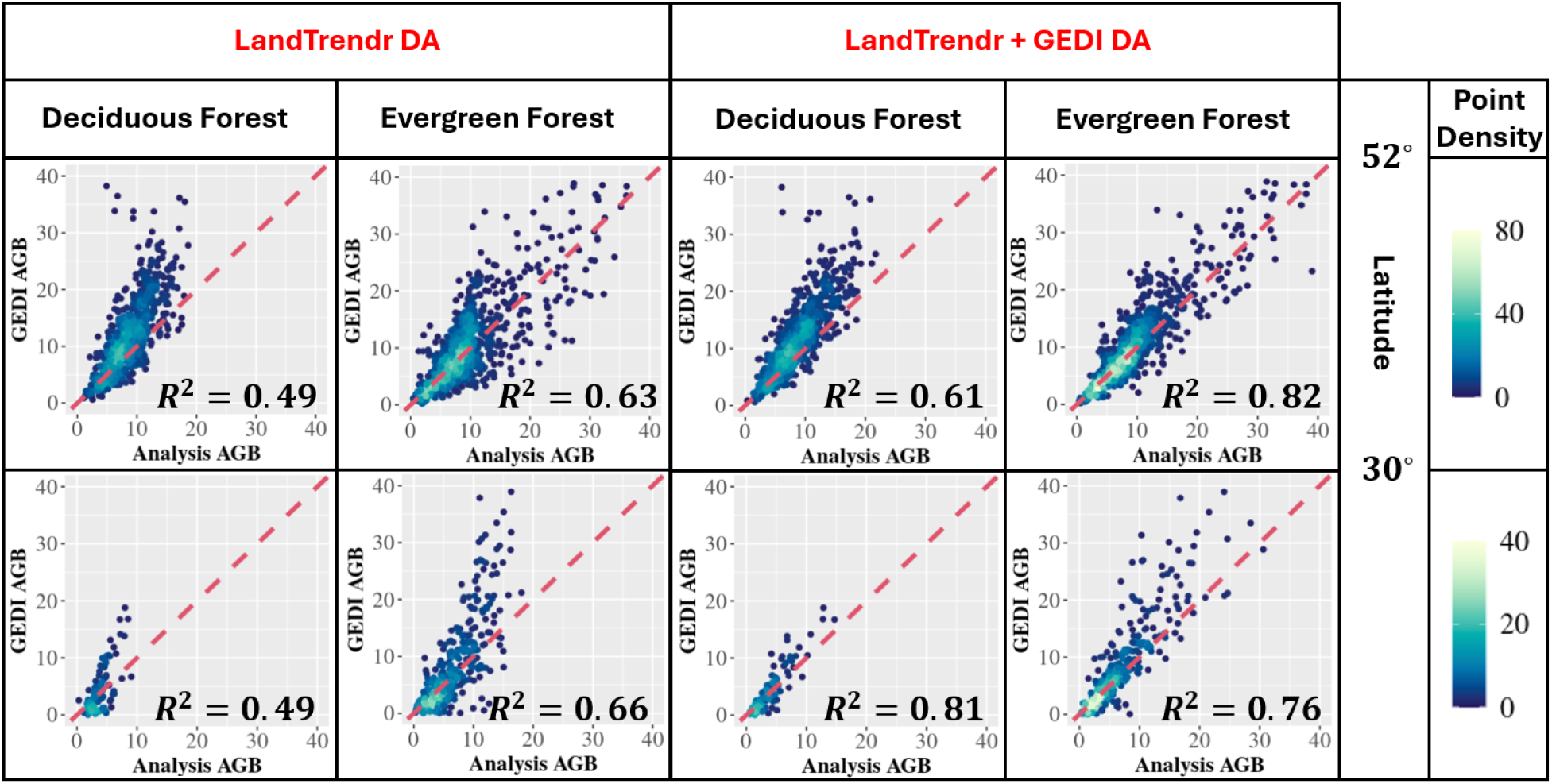
Comparison of the full run and LandTrendr DA results between GEDI and DA AGB split by latitude range (30 - 50 degrees and < 30 degrees) and forest types (deciduous and evergreen forest, respectively), where the red dashed lines are 1:1 lines. The plots are colored from blue to yellow, indicating low to high point density.

In the spatial comparisons, Figure 6 shows the changes in residual errors for both baseline and full run DA relative to the GEDI L4B observation. From the figure, we observed a substantial improvement in accuracy across North America, with the largest improvements in the high biomass forests of the Pacific Northwest and Appalachians, regions where earlier versions of the DA had been systematically underestimating biomass because of saturation issues with LandTrendr. There was also a noticeable improvement over the northern Great Plains. Areas where accuracy decreased were generally diffuse and modest, with the most notable pockets in the northern Rockies and the eastern Maritime Peninsula. Similarly, the AGB uncertainty was notably reduced across CONUS and southern Canada. Across Mexico and Central America, where GEDI footprint density is notably lower, uncertainty reductions were generally smaller in magnitude with occasional pockets of increased uncertainty (Figure 6b).

**Figure 6.**
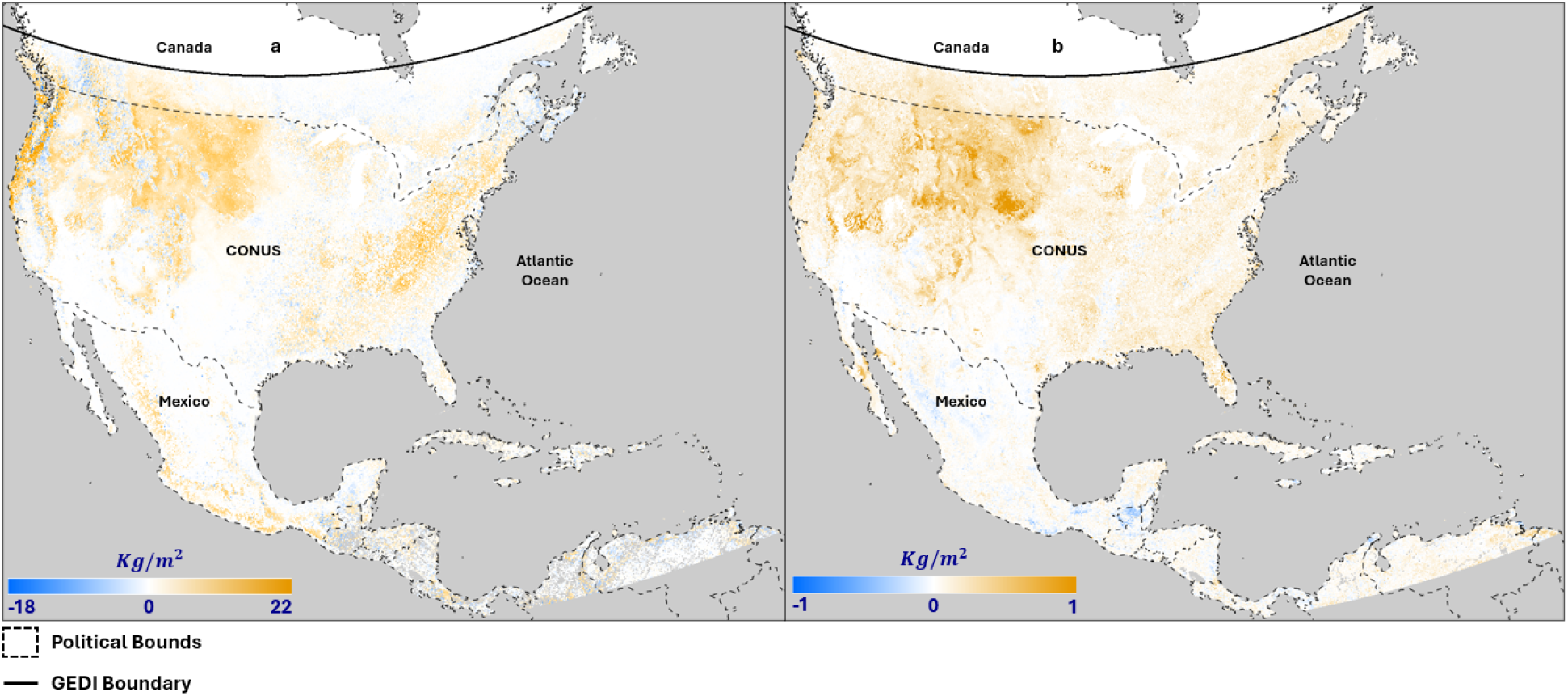
The gridded map of the relative change between the full-run and baseline-run regarding (a) the residual error (a) and (b) predictive uncertainty (b). Yellow means reduced errors, blue means increased errors, grey areas are the ocean. Here, we first calculated the absolute AGB error between the two experiments and the GEDI L4B observation, and then calculated the error reduction between the two experiments for each pixel. The uncertainty change was calculated similarly for the two experiments.

Figure 7 shows that the AGB uncertainty in the baseline-run results decreases continuously over time, whereas the full-run uncertainty is substantially lower and more stable. When we zoom in on the full-run trend, Figure 4.5b shows that the temporal uncertainty for the full-run continues to decrease from 2012 to 2017, 2019 to 2022, and 2023 to 2024. The small increases in uncertainty are due to a sequence of changes in the corresponding observations: 1) LandTrendr discontinuity in 2018; 2) start of GEDI assimilation in 2019; 3) GEDI pause in 2023.

**Figure 7.**
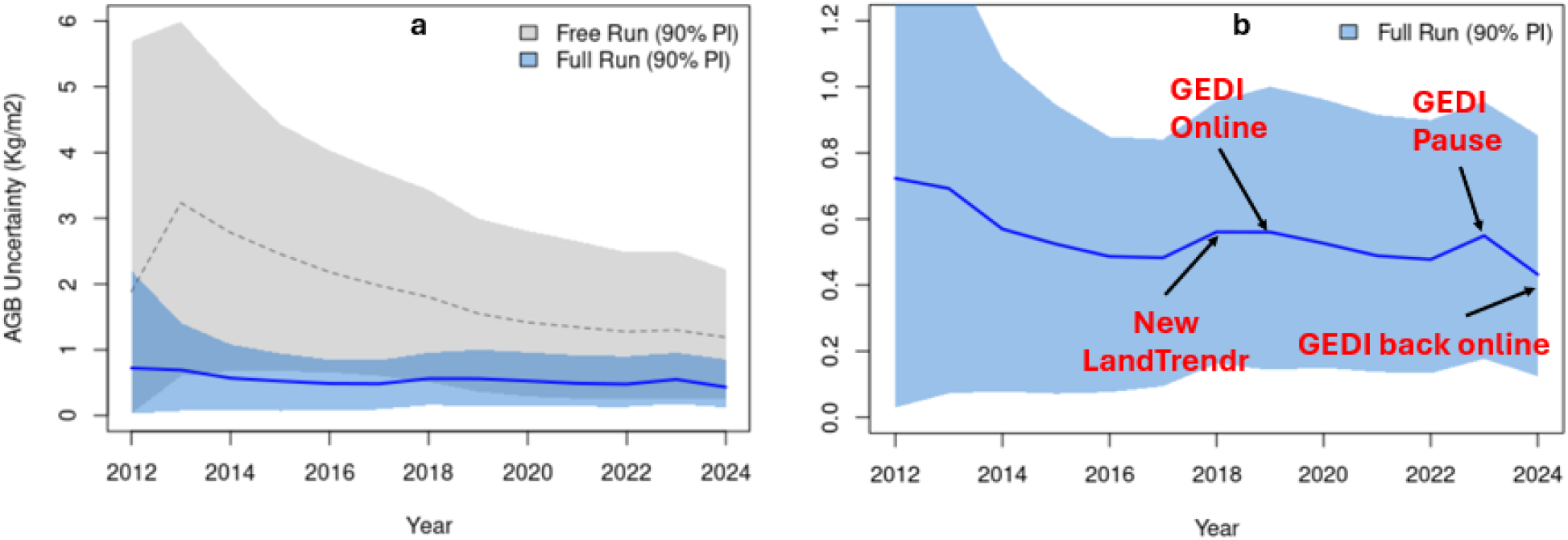
The temporal trends of the predictive error of AGB between the full-run and baseline-run (a), and a zoomed-in full-run trend (b) with specific events labeled on the figure. Here, we calculated the uncertainty using the averaged ensemble standard deviation across the pre-selected locations within the GEDI boundary.

### 3.2. Spatial Error Patterns

Although we achieved substantial improvements in both residual and predictive AGB errors across time and space, we still observed nontrivial areas where the full run performed worse relative to the baseline run. As shown in Figures 8a-b, both the absolute residual error and posterior uncertainty were negatively correlated with the log of the number of high-quality GEDI waveforms, meaning that more GEDI observations lead to lower error. The AGB absolute error was more sensitive to the number of GEDI tiles than the AGB predictive uncertainty, presumably because the AGB uncertainty is affected not only by data quality (the more GEDI observations, the lower the data uncertainty) but also by the AGB pool size, as shown in Supplement 4.8, where about 83% of the pixels with increased uncertainty showed an increase in AGB relative to the baseline-run results.

**Figure 8.**
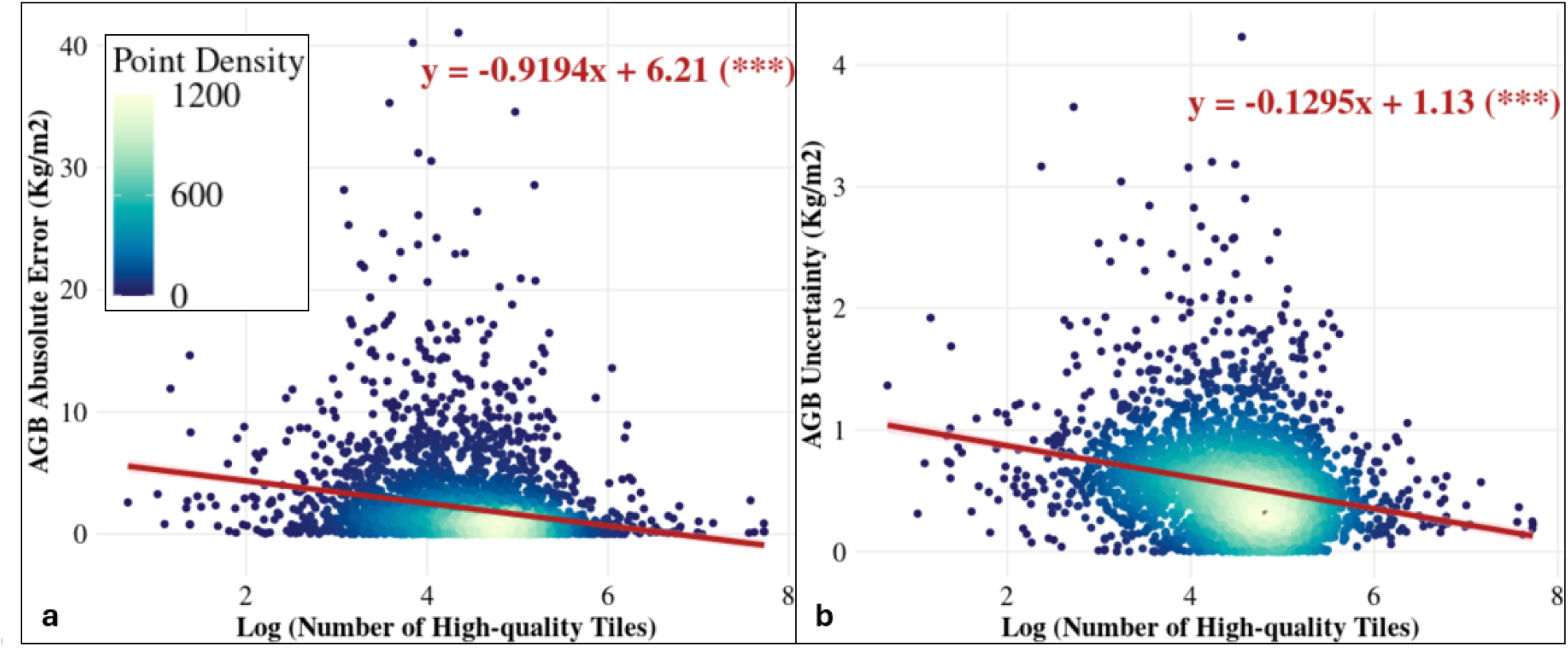
The correlations between the log number of high-quality GEDI tiles and the residual error (a) and predictive error (b). Here, we extracted the number of GEDI tiles for each pre-selected location and then visualized it against the corresponding residual and predictive errors.

Figure 9 visualizes the spatial correlation between AGB predictive skill (residual error and uncertainty) and potential driving factors (elevation, number of GEDI tracks, and annual cloudy days) using local binary Moran’s I plots. Low accuracy is spatially correlated with a low number of GEDI waveforms (Low-low = dark blue) in the Subtropical, the South-Central United States, and parts of the northeastern US (Figure 9a). Reduced AGB precision in Mexico is strongly correlated with lower GEDI waveform counts (Low-low = dark blue) (Figure 9b). Figures 9c-d analyze the spatial correlation among accuracy, elevation, and cloud cover. From the figures, we can see that areas with increased residual error are mostly correlated with high elevation in the western mountainous region. Figure S6 also shows that locations with reduced accuracy are at higher elevations than those with increased accuracy (t = -136.74, df = 84702, p-value < 2.2e-16). High cloud coverage also explains reduced accuracy, particularly in the northeast, and areas with reduced accuracy typically experience higher cloud coverage than those with increased accuracy (t = -114.18, df = 20634, p-value < 2.2e-16).

**Figure 9.**
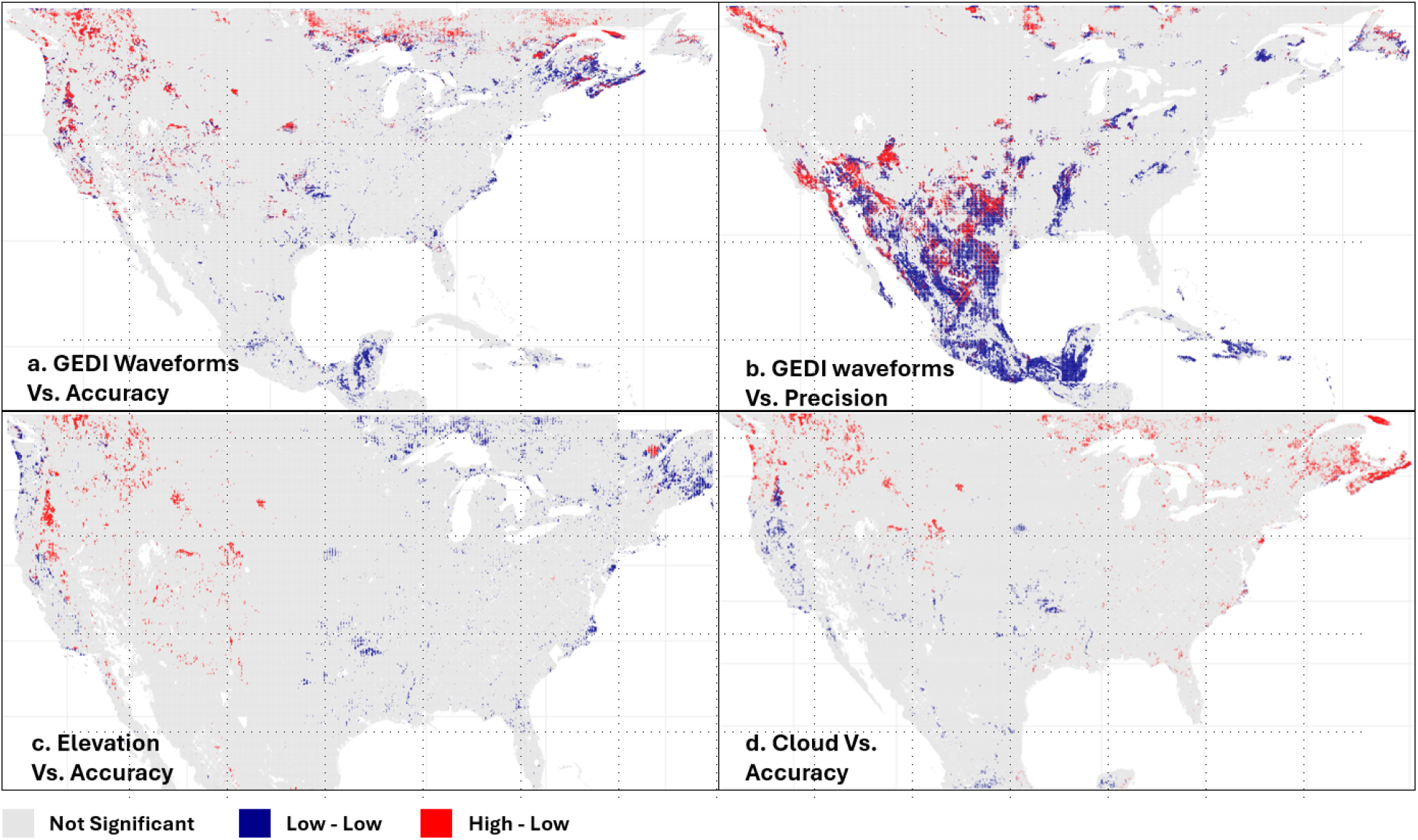
The spatial correlation between a) the number of GEDI waveforms and accuracy; b) the number of GEDI waveforms and the AGB uncertainty reduction; c) elevation and accuracy; d) the annual cloudy days and accuracy. All colors (except grey) show significant spatial correlation (p-value < 0.01) across three clusters between labeled variables shown in each figure.

## 4. Discussion

In this study, we used DA to combine AGB observations from both optical and LiDAR remote sensing. The results show the feasibility of the proposed framework, with low residual and predictive errors across both sets of observations. In addition, errors during validation reveal interesting spatiotemporal patterns that will be used to address our earlier research questions:

### 4.1. What are the spatiotemporal patterns for the residual and predictive errors relative to the baseline model and to the results of the previous North American DA work?

Specifically, assimilating both GEDI and LandTrendr AGB yields low residual errors relative to the corresponding data constraints (*R*^2^ from 0.64 to 0.86; RMSE from 5.61 to 2.69 *kg*/*m*^2^) and low uncertainty relative to the baseline-run results (from 1.89 to 0.72 *kg*/*m*^2^ in 2012 and from 1.19 to 0.43 *kg*/*m*^2^ in 2024). Beyond that, the full-run results consistently outperformed the LandTrendr-only DA results, with *R*^2^ improvements from 0.1 to 0.32.

By harmonizing information from both optical and LiDAR remote sensing, we are ultimately able to borrow strength from both sensors with low errors in space and time. This example of using DA to reconcile multiple observations of the same variable opens the door to broader applications involving additional sources of biomass information (e.g., inventory, Lichstein et al., 2014; radar, Liu et al., 2026) and extending this to using multiple sources of information to constrain other state variables (e.g., LAI, soil moisture) (Samadzadegan et al., 2025).

As Figure 4 shows, LandTrendr accuracy begins to decline after GEDI appears, while GEDI accuracy increases from 2019 to 2024 (except in 2023, when GEDI paused). This reveals a key advantage of our framework: the ability to objectively weight multiple data constraints based on their quantified uncertainties, allowing the system to dynamically prioritize the most reliable product. Beyond that, prioritizing GEDI highlights the advantage of LiDAR AGB estimates over the optical remote sensing product, so the annualized GEDI footprint aggregations receive more weight in the DA workflow, which also demonstrates the reliability of our GEDI aggregation workflow.

While GEDI provides substantial improvements over both the baseline and previous DA experiments, optical remote sensing estimates (i.e., LandTrendr AGB) also provide comprehensive coverage of AGB across space and time, especially when there are no GEDI measurements. LandTrendr can be accurate across low-to-medium biomass regions and provides complete coverage over CONUS at 30m gridded pixels, addressing the GEDI platform’s data sparsity. Together, combining both LiDAR and optical AGB estimates in our DA workflow provides an accurate C MRV platform that can borrow strength from multiple remote sensing platforms.

Another benefit of using aggregated GEDI footprints is informing disturbance, because relative to optical remote sensing platforms, LiDAR can more easily penetrate the canopy and provide accurate measurements of forest 3-D structure. LiDAR observations can detect subtle changes in forest structure, providing a powerful tool for monitoring forest disturbance (Vallet & Mouillo, 2026). The GEDI aggregation proposed in this study can provide annual LiDAR aggregations, unlike GEDI L4B products, which average across the entire GEDI history. Thus, it can enable both forest disturbance detection and recovery monitoring at a finer temporal frequency.

### 4.2. What are the driving factors that affect both the residual and predictive errors?

Although we achieved reduced errors across most of space and time, we still observed areas with increased AGB residual errors in the Subtropical, the South-Central United States, and parts of the northeastern US, and increased AGB uncertainty in the Subtropical. Our results suggest strong correlations between GEDI waveform counts, accuracy, and precision (Figures 8 a-b), and between the AGB pool size and uncertainty (Supplement 8). Furthermore, areas of increased residual error in AGB appear to reflect the combined impacts of GEDI data quantity, elevation, and cloud cover. Specifically, the drivers of high uncertainty appear to be high elevation across the northern Rockies, cloud cover in the northeast in general (and Maritime Provinces specifically), and reduced GEDI data volume in Mexico (Figure 9).

In regions with high cloud cover, adding AGB estimates from radar/microwave sensors could greatly reduce errors in AGB estimates (Padalia et al., 2023), as these signals readily penetrate clouds. Beyond that, while the LiDAR platform penetrates the canopy better than optical remote sensing, it will still be limited in high-biomass vegetation. Therefore, we will consider combining LiDAR with microwave and optical remote sensing platforms (Shendryk et al., 2022), since microwave remote sensing can penetrate both dense clouds and forests and improve AGB estimation accuracy. Finally, an important future direction is to better account for the identified sources of uncertainty (e.g., elevation, cloud cover, and GEDI waveform counts) as additional covariates during the ML debiasing process.

## 5. Conclusion

This study presents a novel application of DA to harmonize AGB estimates from both optical and LiDAR platforms, substantially reducing residual and predictive errors across space and time. As a proof-of-concept, this analysis illustrates the value of combining GEDI footprints at annual temporal resolutions and the ability of satellite LiDAR to help correct biases in optical AGB estimates, particularly in high biomass regions. In addition, the sources of AGB residual error and uncertainty are strongly correlated with elevation, cloud coverage, and GEDI data density, though which effect dominates differs substantially across space.

## Supporting information

https://docs.google.com/document/d/1qMdWb1QhYOH3OrILehL2VAThlXK87Ws9yxiWo5oTf_Q/edit?usp=sharing

## 6. Acknowledgment

This work is supported by NASA CMS Grant: 20-CMS20-0030 and NSF MSB Grant: 2406258. We would also like to thank Emily Beasley for her generous feedback on the content, style, and organization of this paper.

