## Supplementary material for "Harmonizing Repeated Multi-Platform Optical and LiDAR Observations for Continental-Scale Terrestrial Carbon Cycle Data Assimilation": https://docs.google.com/document/d/1qMdWb1QhYOH3OrILehL2VAThlXK87Ws9yxiWo5oTf_Q/edit?usp=sharing

**Supplements.**

**Supplement 1. Site Selection Algorithm.**

The site-selection algorithm was developed to select representative locations across eco-climatic variations, conditional on each vegetation type from the MODIS PFT land cover classifications. In this step (Figure S1), we first compiled the locations of ~400 "anchor" sites, defined as locations with ground-based networks (e.g., AmeriFlux, MexFlux, NEON, LTER, LTAR, ForestGEO, CZO, PhenoCam) that are of scientific interest, and combined this with the ~500 sites from our previous CONUS SDA (Dokoohaki et al., 2022), which were selected using an algorithm analogous to that described below.

Next, we calculated the number of locations for each land cover class proportionally to its spatial coverage, with a minimum of 200 sites per class (Figure S1), totaling 8,000 sites, to balance computational demand and representation of eco-climatic gradients. The remaining locations, excluding the anchor sites, were weighted for sampling to aid in future FIA and GEDI validations. The weighting was conducted under the CONUS Albers projection, which preserves area across latitudes, and the weight layer was calculated as the number of by-pixel FIA plots and high-quality GEDI waveforms (Table S1). Additionally, because the GEDI orbit stops at 52°N, samples for each land cover class were further split along this latitude proportionally to area.

We then assembled a suite of environmental covariates selected to explain variability in carbon pools and fluxes (Table S1). A global forest age map circa 2010 (Besnard et al., 2021) was used to represent the time elapsed since the most recent disturbance, adjusted to 2024, and supplemented with the MODIS land cover product to account for post-2010 land cover changes and the age of non-forest land cover classes. Additional covariates include the global AGB in 2010, annual median maps for MODIS LAI and SMAP root zone SM, elevation from the STRM digital elevation model dataset (Jarvis et al., 2008), and soil properties (e.g., organic carbon content (ORC), pH, nitrogen, and sand proportion) from the SoilGrids database (Hengl et al., 2017). Climatic covariates, including temperature, precipitation, solar radiation, and vapor pressure, were generated based on ERA5 annual mean estimates. For clustering, all data layers were normalized to a 0–1 scale based on their min/max values, except land cover, which was converted to a factor.

Then, we employed the Hierarchical k-means clustering method (Alboukadel & Fabian, 2020) to identify unique clusters for each land cover class, using 60,000 weighted sampled locations per class to accommodate the algorithm's memory limitations. The k-means clustering was executed based on the maximum total within-cluster sum of squares (WCSS) obtained from the elbow point on a plot of WCSS versus cluster sizes from 1 to 20. After dividing the number of locations to be sampled evenly among clusters, we removed the anchor sites and sampled from each cluster in proportion to the FIA and GEDI weight layer.

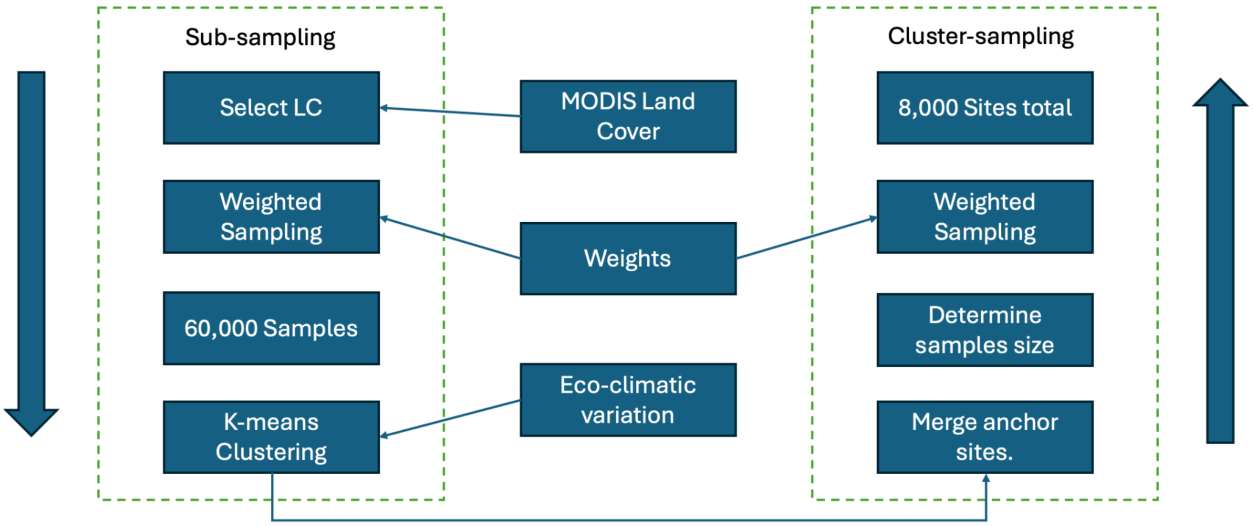


*Figure S1. Site selection workflow. We first generate 60,000 random subsamples per MODIS land cover type using weighted sampling, prioritizing FIA and GEDI coverages. Next, we generate k-means clusters based on the eco-climatic variables (Table S1), exclude the pre-selected sites (e.g., NEON, AmeriFlux, FluxNet), and determine the optimal cluster size by the elbow location in the WCSS curve. Finally, weighted sampling is applied to each cluster.*

*Table S1. Environmental covariates used for k-means clustering, debiasing, and ML emulators. [1]Forest age 2010 base map updated annually using MODIS land cover and extended to non-forest pixels.*

| Data Source | Temporal Domain | Spatial Resolution | Temporal Frequency | DOI |
|---|---|---|---|---|
| ERA5: temperature, precipitation, solar radiation, vapor pressure | 2012-2024 | 30 km | annual | 10.24381/cds.adbb2d47 |
| STRM DEM | 2000 | 90 m | static | https://srtm.csi.cgiar.org. |
| Forest Age | 2010 | 1 km | annualized[1] | https://doi.org/10.17871/ForestAgeBGI.2021 |
| GEDI level 4B: number of samples | 2019-04-18 to 2023-03-16 | 1 km | static | 10.3334/ORNLDAAC/2299 |
| MODIS MCD12Q1 Land Cover | 2001 to 2024 | 500 m | annual | 10.5067/MODIS/MCD12Q1.061 |

| | | | | |
|---|---|---|---|---|
| AGB 2010 | 2010 | 300 m | static | 10.3334/ORNLDAAC/1763 |
| SoilGrids: ORC, pH, nitrogen, and sand | NA | 250 m | static | https://doi.org/10.5194/soil-7-217-2021 |
| FIA Plot Density | 2000 to 2024 | NA | static | https://doi.org/10.2737/RDS-2001-FIADB |

**Supplement 2. Equations for the Multi-observational Tobit-Gamma Ensemble Kalman Filter (MoTGEnF).**

**1. Prior for Process Precision (Q).**

The code loops over $N$ state variables, assigning a Gamma prior to the precision (the inverse of variance) for each variable:

$$Q_i \sim Gamma(aq_i,\ bq_i)\ for\ i = 1, ..., N$$

- $Q_i$: The precision of the process error for state variable $i$.
- $aq_i$: The shape parameter of the Gamma distribution.
- $bq_i$: The rate parameter of the Gamma distribution.

**2. Latent State (Process) Model.**

The latent state variable $X_i$ is normally distributed around the modeled deterministic expectation ($X.mod_i$):

$$X_i \sim \mathcal{N}(X.mod_i, Q_i)\ for\ i = 1, ..., N$$

- $X_i$: The true latent state.
- $X.mod_i$: The expected state (likely the output of a forward deterministic model).
- $Q_i$: The process variance.

**3. Observation Model.**

The observation model maps the observations ($Y_j$) to the corresponding latent states ($X_i$) using an index/mapping vector $H$:

$$Y_j \sim \mathcal{N}(X_{H(j)}, r_{j,j})\ for\ i = 1, ..., YN$$

- $Y_j$: The $j^{th}$ observation.
- $X_{H(j)}$: The specific latent state corresponding to observation $j$ (where $H(j)$ is the mapping index).
- $r_{j,j}$: The observation variance for the $j^{th}$ observation, taken from the diagonal of an observation precision matrix $r$.

**Supplement 3. SIPNET.**

We used the Simplified Photosynthesis and Evapotranspiration Model (SIPNET) (https://github.com/PecanProject/sipnet), a simple pool-based process model, as the scaffold for making forward predictions of all C pools and fluxes and reconciling predictions with observations (Braswell et al., 2005). SIPNET was selected based on its low computational cost and its simplified process representations of coupled carbon and water cycling, which facilitate interpretability and data assimilation.

SIPNET contains four vegetation carbon pools (leaf, stem, coarse roots, and fine roots) and one soil carbon pool (Figure 5). Photosynthesis is calculated using a light-use-efficiency approach that is a function of maximum photosynthesis, LAI, and photosynthetically active radiation, with modifiers for air temperature, vapor pressure deficit, and soil moisture. SIPNET also has a simple representation of vegetation C allocation based on fixed fractions to each plant carbon pool. For deciduous systems, SIPNET was run in “prescribed phenology” mode based on the day-of-year of leaf-on and leaf-off. Losses from plant carbon pools are based on fixed turnover rates and maintenance respiration, which is a Q10 function of temperature. Soil carbon is increased by wood and leaf litter and reduced by soil respiration. Finally, soil moisture is modeled using a simple bucket model approach, with precipitation as input and evaporation, transportation, and drainage as outputs. To match the assimilated observations, SOC was modeled down to a 2m depth, while soil moisture was modeled to a 1m depth.

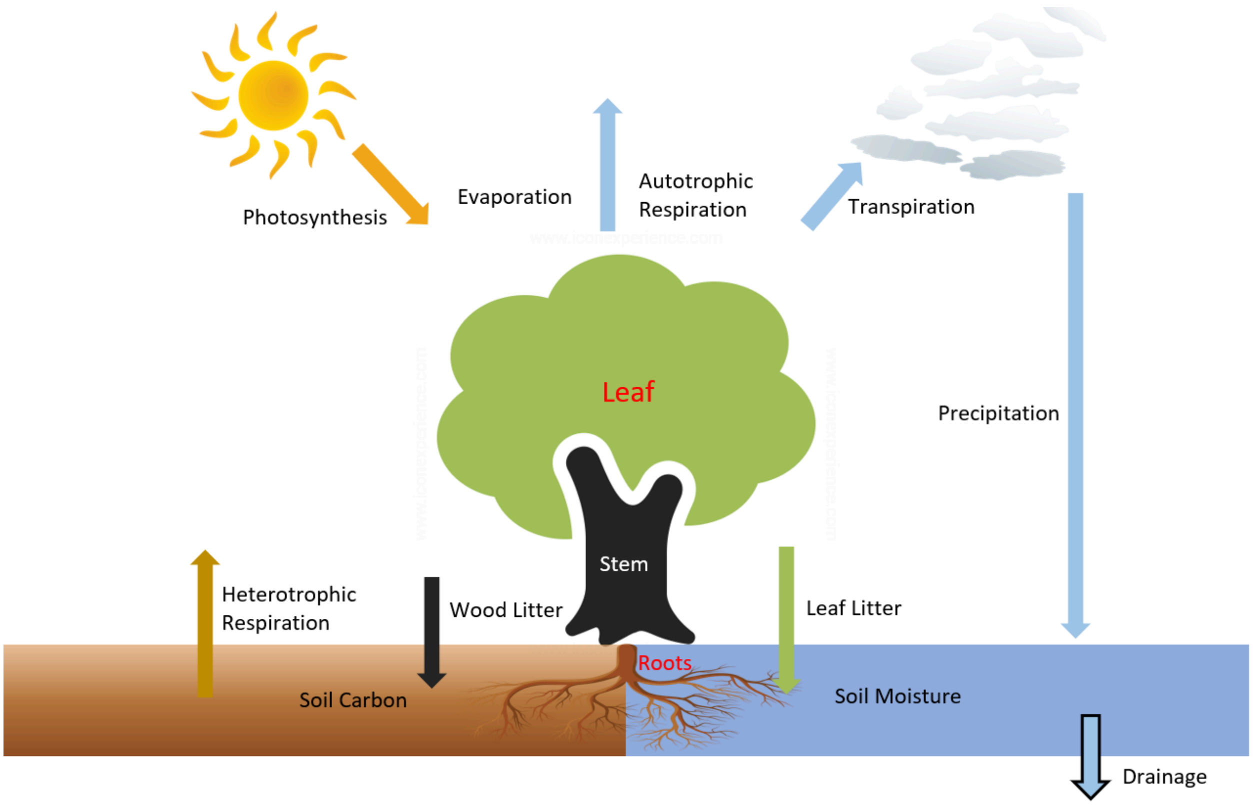


*Figure S2. Flowchart of the SIPNET model. The model includes four vegetation carbon pools (leaf, stem, coarse roots, fine roots) and one soil carbon pool. Photosynthesis adds carbon to the system, which is then allocated to the four vegetation pools. Meanwhile, autotrophic and heterotrophic respiration remove carbon from the vegetation and soil, respectively. Vegetation pool turnover (a.k.a. litter) adds to soil carbon, and precipitation adds soil moisture, the latter of which is removed by transpiration, evaporation, and drainage.*

**Supplement 4. Data Preparations.**

**4.1. Initial Conditions (ICs)**

The initial conditions (ICs) define the starting states of the ecological model. To initialize LAI, we used the maximum MODIS LAI observations from 2011-06-01 to 2011-08-31 after removing outliers (>0.95 or <0.05 quantiles), with QA/QC values of either "000" or "001". To avoid singularity issues in Bayesian analyses, we set a minimum of 0.66 standard deviations (SD) for LAI (Viskari et al., 2015). The AGB initial conditions were based on a 2010 global AGB map (Spawn et al., 2020). SM was initialized from meteorological reanalysis (Dorigo et al., 2019). We eventually generated 100 ensembles for each IC and location using the reported mean and SD for each IC. All ICs were sampled using the zero-truncated normal sampling (Olaf et al., 2018) independently for each location.

SOC was initialized using the ISCN database (Nave et al., 2016) with less than 200 cm of soil depth. To reduce the potential double use of data in both initialization and SDA between ISCN and SoilGrids, SOC priors were generated by sampling ISCN measurements with respect to each of the fifteen EPA level 1 ecoregions (Omernik, 1995). The global mean and SD will be used for places outside of the EPA map.

*Table S2. Data used as model inputs. Here, we list the data source, temporal domain, and frequency, as well as the spatial resolution and data DOI for model drivers (ERA5 meteorology, MODIS phenology), initial conditions (e.g., AGB, LAI, SM, and SOC), and soil physical variables.*

| Data Source | Temporal Domain | Spatial Resolution | Temporal Frequency | DOI |
|---|---|---|---|---|
| ERA5 | 2012 to 2024 | 30 km | 3-hour | 10.24381/cds.adbb2d47 |
| MODIS Phenology | 2012 to 2024 | 500 m | 1-year | 10.5067/MODIS/MCD12Q2.061 |
| AGB 2010 | 2010 | 300 m | NA | 10.3334/ORNLDAAC/1763 |
| MODIS LAI | 2011 | 500 m | 8-day | 10.5067/MODIS/MOD15A3H.006 |
| Gridded Soil Moisture | 2011 | 0.25 degree | 10-day | 10.24381/cds.d7782f18 |
| ISCN SOC | 2012 to 2014 | NA | NA | 10.17040/ISCN/1305039 |
| SoilGrids soil texture data | NA | 250 m | static | 10.5194/soil-7-217-2021 |

**4.2. Model Inputs.**

Model parameter ensembles were sampled from by-PFT joint posterior parameter distributions generated through previous calibrations using a Hierarchical Parameter Data Assimilation (HPDA) approach against carbon and water flux measurements from 22 AmeriFlux towers (Dokoohaki et al., 2022; Fer et al., 2018, 2021). Phenological drivers, including leaf-on and leaf-off dates, were extracted from the MODIS Land Cover Dynamics (Friedl et al., 2022) database, which provides annual global land surface phenology metrics from 2001 to the present at a 500 m spatial resolution. We used the “mid-green-up” and “mid-green-down” bands when EVI2 (enhanced vegetation index 2) crossed 50% of the seasonal amplitude to identify leaf-on and leaf-off dates. Ensembles of site-specific soil texture parameters (e.g., soil water-holding capacity & drainage rate) were generated based on the SoilGrids dataset (Supplement 5) using the look-up tables and pedotransfer functions in Cosby et al. (1984). The ERA5 reanalysis was used as meteorological drivers (Hersbach et al., 2023), given its continuous global coverage with ten ensembles, which were randomly assigned to the 100 model ensembles.

### 4.3. Observational Data Constraints

This study simultaneously assimilated AGB, LAI, SM, and SOC, building on our previous work assimilating the same constraints across CONUS NEON sites (Zhang et al, 2025). We used the LandTrendr product (Hooper & Kennedy, 2018) as the AGB constraint, which uses LandTrendr estimates of the Landsat composite within CONUS, calibrated against forest inventory data, providing AGB means and uncertainty estimates until 2017 and only AGB means from 2018 to 2023. To predict uncertainties for 2018 to 2023, we trained the Random Forest algorithm against the reported uncertainties from 2012 to 2017 using covariates from the site-selection algorithm (Supplement 1).

MODIS LAI observations were extracted from June 1st to August 31st for each year (with the QA/QC values falling within "000" or "001") and the 0.025-0.975 quantiles to avoid noisy estimates due to cloud cover. We then selected the estimates that are closest to July 15 for each year and location, with 90% of the values falling within 8 days of the target date (see Figure S3). Similar to LAI IC, we assigned a minimum SD of 0.66 to the extracted LAI observations.

The SM is constrained by the SMAP Level 4 root-zone SM estimates, which use a data assimilation system to merge L-band brightness temperature observations with the NASA Catchment land surface model (Reichle et al., 2023). The estimates provide continuous SM to 100 cm depth (the same root-zone definition as SIPNET) with associated uncertainties. In this study, we extracted SMAP estimates at midnight on July 15 each year from 2015 to 2024.

SOC estimates were obtained to a depth of 200cm from the OCD band of SoilGrids data. Since uncertainty estimates are given as a mean and three quantiles (0.05, 0.5, 0.95) for each soil layer, we converted these into an overall column-integrated mean and standard deviation by fitting Gamma distributions across six soil depths (0 - 5cm, 5 - 15cm, 15 - 30cm, 30 - 60cm, 60 - 100cm, and 100 - 200cm) and aggregating the distribution's mean and variance through the entire soil profile.

*Table S3. Details for the data used as observations. Here, we summarize the data details by name, temporal domain, frequency, spatial resolution, and data DOI.*

| Data Source | Temporal Domain | Spatial Resolution | Temporal Frequency | DOI |
|---|---|---|---|---|
| LandTrendr AGB | 2012 to 2023 | 30 m | 1-year | 10.1088/1748-9326/aa9d9e |
| MODIS LAI | 2012 to 2024 | 500 m | 8-day | http://doi.org/10.5067/MODIS/MOD15A3H.006 |
| SMAP level 4 root zone soil moisture | 2015 to 2024 | 9 km | 3-hour | doi:10.5067/02LGW4DGJYRX |
| SoilGrids OCD | NA | 250 m | NA | https://doi.org/10.5194/soil-7-217-2021 |

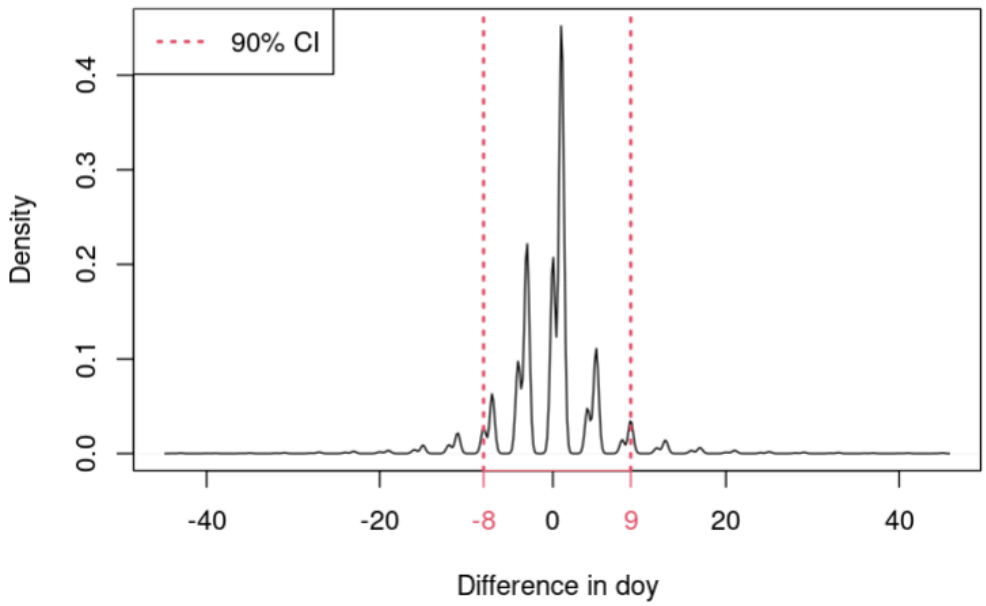


*Figure S3. The differences in the day of the year of our LAI selection date relative to July 15th for each year and site.*

**Supplement 5. The workflow of generations for soil water holding capacity and drainage rate.**

The soil moisture used in the model is limited by the soil water holding capacity. These critical parameters and the drainage rate were estimated through a multi-step process:

- Soil Texture Data: We used the mean and quantiles (5th, 50th, and 95th) of sand, clay, and silt fraction data from SoilGrids (Hengl et al., 2017) for each site and depth.
- Ensemble Generation: These data were used to fit Dirichlet distributions using the moment matching method, which fits the quantiles with means as known moments. Then, the 100-ensemble soil texture datasets were generated for each site and depth.
- Parameter Inference: Soil physical properties, including water-holding capacity and drainage rate, were inferred for each ensemble member using pedotransfer equations (Cosby et al., 1984).
- Integration: The final soil parameters were integrated over the top 100 cm to align with the depth of the SMAP soil moisture data for each site.

**Supplement 6. Machine Learning Debias workflow.**

Systematic errors in model forecasts can persist across cycles and reduce the skill of the data assimilation. To address this, we developed a machine learning (ML) framework that corrects ensemble forecasts before they are passed into the filter. The goal is to shift the ensemble mean to remove bias while leaving the ensemble spread intact, so that uncertainty propagation remains governed by the filter.

At each assimilation cycle, the process-based model (SIPNET) generates an ensemble of forecasts for each state variable. From this ensemble, we compute the mean forecast, denoted $\mu^f$, and the ensemble covariance, denoted $\Sigma^f$. Given an observation $y^{obs}$, we define the residual as $\varepsilon = y^{obs} - \mu^f$. These residuals are then merged with a set of environmental covariates, including climate, soils, land cover, disturbance history, and topography. In addition, the forecast itself was explicitly included as a predictor, since systematic errors often scale with forecast magnitude.

The debiasing model is a stacked regression framework implemented in Python and called from R via the reticulate interface. For each state variable, we fit two base learners: a K-Nearest Neighbors regressor (with the number of neighbors k selected by cross-validation) and an ExtraTrees ensemble. Both models are trained to map the combined predictor set (covariates plus forecast) to the observed residuals. Their predictions are then blended with a weight w between 0 and 1, chosen to minimize root mean squared error on the training set. The resulting blended model produces a predicted residual $\hat{\varepsilon}$ for the next timestep.

To apply the correction, we add the same offset $\hat{\varepsilon}$ to each ensemble member. In other words, the corrected ensemble is given by $y^{f,corr} = y^f + \hat{\varepsilon}$. This procedure shifts the ensemble mean ($\mu^{f,corr} = \mu^f + \hat{\varepsilon}$) while leaving the covariance unchanged ($\Sigma^{f,corr} = \Sigma^f$). By construction, the debiasing step only adjusts the location of the ensemble distribution, ensuring that spread and shape are preserved for the subsequent assimilation update.

These bias-reduced forecasts are then passed to the Tobit-Gamma Ensemble Filter (TGEnF) for assimilation with observations. Cross-validation confirmed that including the forecast itself as a predictor significantly improved residual prediction, and feature importance analysis showed that the forecast was consistently the single most important predictor across all variables. Environmental covariates such as disturbance history, precipitation, temperature, and soil texture provided additional explanatory power, but their effects were secondary to the forecast itself. Together, these results show that the ML debiasing framework reduces systematic forecast–observation mismatches and stabilizes the assimilation process by aligning the prior mean with observational evidence before the filter update.

**Supplement 7. The discontinuity of the LandTrendr AGB estimates at year 2017 across 8,000 locations.**

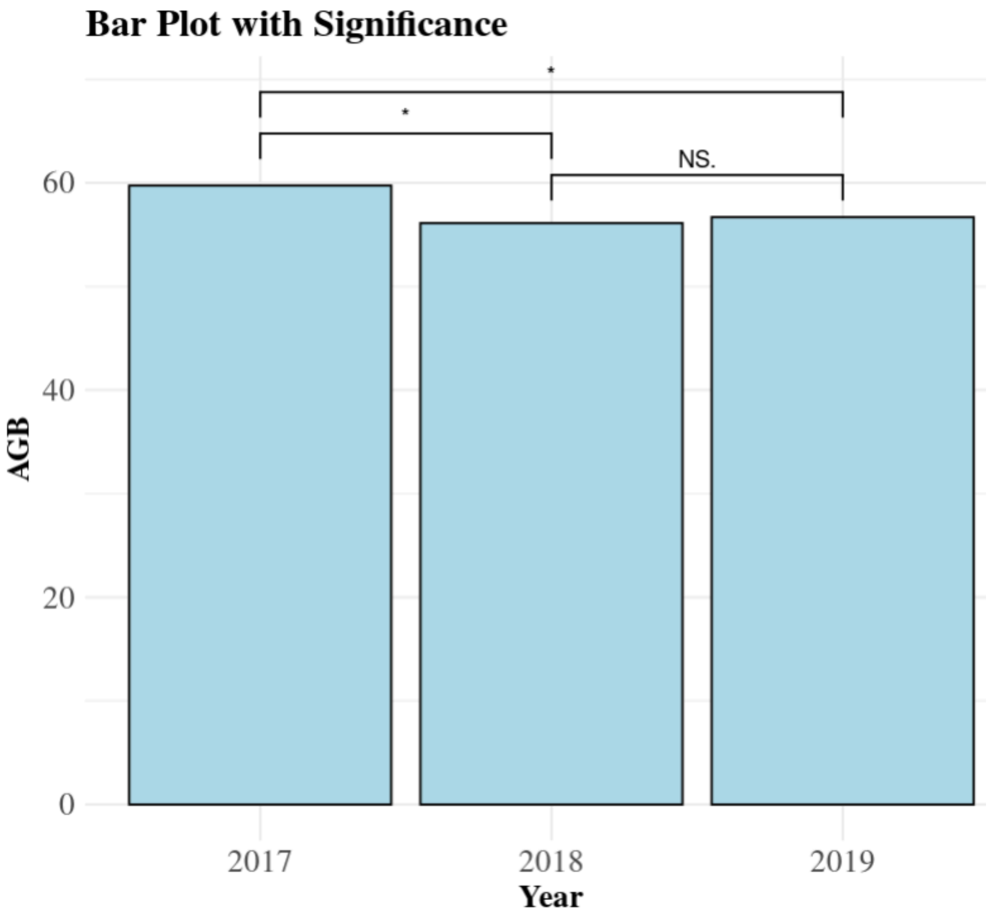


*Figure S4. Bar plot of LandTrendr AGB observations across selected 8,000 locations from 2017 to 2019. From this figure, we can see that the LandTrendr AGB is statistically significantly reduced after 2017.*

**Supplement 8. Density distribution of AGB change between full-run and baseline-run across pixels that show increased uncertainty.**

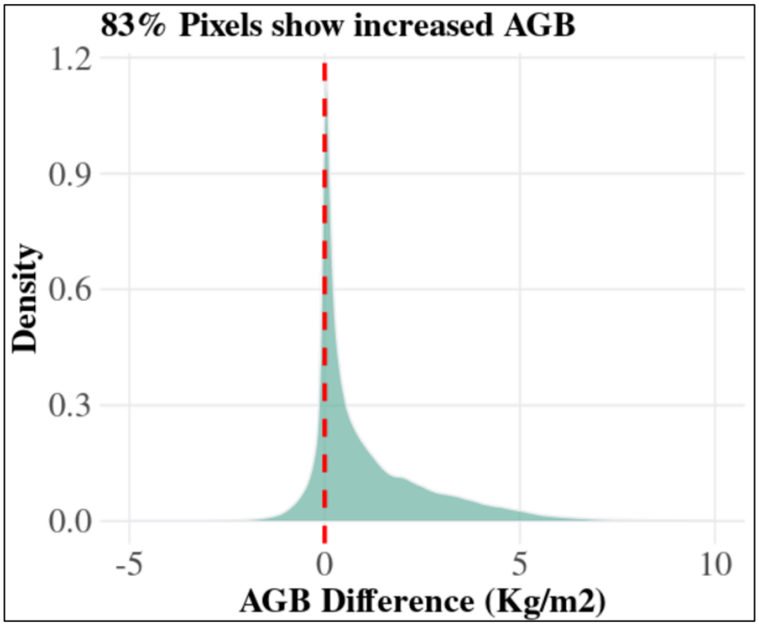


*Figure S5. The density distribution of AGB difference when subtracting the baseline AGB estimates from the full-run AGB estimates.*

**Supplement 8.**

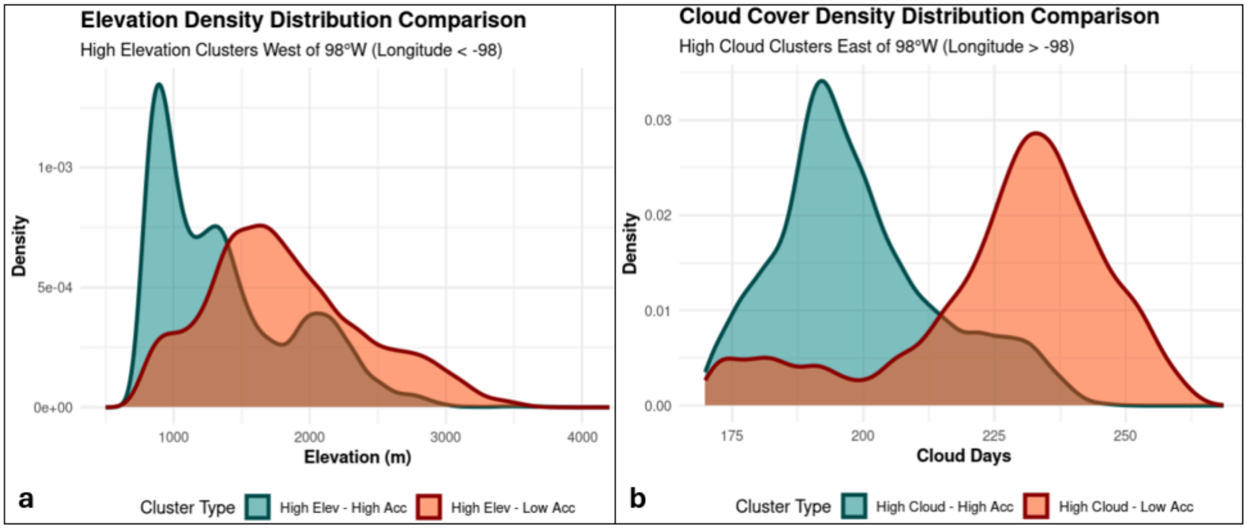


*Figure S6. The density distributions of a) elevation and b) cloudy days between the improved and worsened accuracy clusters.*
